# DELLA Proteins Recruit the Mediator Complex Subunit MED15 to Co-activate Transcription in Land Plants

**DOI:** 10.1101/2023.11.01.565078

**Authors:** Jorge Hernández-García, Antonio Serrano-Mislata, María Lozano-Quiles, Cristina Úrbez, María A Nohales, Noel Blanco-Touriñán, Huadong Peng, Rodrigo Ledesma-Amaro, Miguel A Blázquez

## Abstract

DELLA proteins are negative regulators of the gibberellin response pathway in angiosperms, acting as central hubs that interact with hundreds of transcription factors and regulators to modulate their activities. While the mechanism of transcription factor sequestration by DELLAs to prevent DNA binding to downstream targets has been extensively documented, the mechanism that allows them to act as co-activators remains to be understood. Here, we demonstrate that DELLAs directly recruit the Mediator complex to specific loci in Arabidopsis, facilitating transcription. This recruitment involves DELLA amino-terminal domain and the conserved MED15 KIX domain. Accordingly, partial loss of MED15 function mainly disrupted processes known to rely on DELLA co-activation capacity; including cytokinin-dependent regulation of meristem function and skotomorphogenic response, gibberellin metabolism feedback, and flavonol production. We have also found that the single DELLA protein in the liverwort *Marchantia polymorpha* is capable of recruiting MpMED15 subunits, contributing to transcriptional co-activation. The conservation of Mediator-dependent transcriptional co-activation by DELLA between Arabidopsis and Marchantia implies that this mechanism is intrinsic to the emergence of DELLA in the last common ancestor of land plants.

**Significance Statement:** DELLA proteins are plant-specific transcriptional hubs integrating environmental signals with endogenous cues. In order to regulate downstream processes, DELLAs modulate the activity of hundreds of transcription factors and transcriptional regulators in various ways. Here, we describe the molecular mechanism underlying DELLA co-activator function. We show that DELLAs act as transcriptional activators in eukaryotic cells by interacting with the Mediator complex subunit MED15. Mediator function is necessary to regulate a subset of DELLA-regulated responses that are mediated by direct co-activation of DELLA-Transcription factors complexes. We further show that this mechanism is present in bryophyte DELLAs, and thus represents a conserved mechanism of DELLA function in land plants.

## Introduction

A key mechanism allowing for optimal use of resources in land plants relies on the ability of DELLA proteins to integrate environmental signals and transduce this information to transcriptional programs (1, 2). This central function results from the combination of two features: the influence of environmental conditions on DELLA protein levels and activity, and DELLA’s extensive capacity to interact with dozens of transcription factors (TFs) and transcriptional regulators (3, 4).

In vascular plants (tracheophytes), environmental regulation of DELLA levels is mostly exerted through gibberellin (GA)-dependent DELLA polyubiquitylation and subsequent proteasomal degradation (5). Perception of GAs by the GID1 receptor promotes the interaction with the N-terminal domain of DELLA proteins and the recruitment of the F-box protein SLEEPY1/GID2. In addition, DELLA levels are regulated under certain environmental conditions by GA-independent COP1-mediated proteasomal degradation, and their activity is further modulated by posttranslational modifications (6, 7). In bryophytes, which lack known GID1 receptors, DELLA levels have been suggested to be primarily controlled through transcriptional regulation (8). In contrast to the proposed differences in DELLA regulation among land plants, the intrinsic property to interact with TFs is conserved in all land plant lineages and likely represents an ancestral function of these (9). Depending on the interacting partner, DELLAs affect transcription through two main alternative mechanisms: they either hinder the binding of TFs to their target DNA sequences (10, 11), or are recruited to the chromatin by TFs, where they can act as transcriptional coactivators (3, 12, 13). The ability of DELLAs to activate transcription has been profusely documented, residing in the N-terminal domain, and overlapping with the GID1-interacting region (14, 15). In fact, the conservation of the GID1-interacting motifs in bryophytes could be linked to the functional conservation of the transcriptional activity.

Despite the relevance of DELLAs as transcriptional coactivators, the actual mechanism by which this activity is established is still unknown. In eukaryotes, selective transcriptional activation by a TF or a TF-coactivator pair commonly entails the recruitment of the Mediator complex (16–18). This complex is comprised of more than 20 subunits organized in three main core modules: head, middle, and tail, alongside an additional regulatory module. The head and middle modules are directly involved in the formation of the RNA polymerase II (RNAP II) pre-initiation complex, while subunits of the tail module facilitate the recruitment of Mediator to specific loci through the interaction with TFs or TF-coactivator complexes (19, 20). Among these subunits, MED15 stands out as a common target of many activators in different species, and it has been shown to mediate transcriptional activation of up to 85% of the activator TFs in yeast (21). A feature of MED15 proteins is the presence of a conserved kinase-inducible domain interacting (KIX) domain (22, 23), a domain involved in the interaction with transcriptional activation domains (TADs). In Arabidopsis, some TFs have been identified as putative interactors of the KIX domain (24), but the relevance of a TF-MED15 pair has been demonstrated only for the transactivation activity of the TF WRINKLED1 (22). Here, we have studied the link between DELLA co-activator function and its transactivation activity. Following predictions of a possible TAD, we demonstrate that DELLA proteins contain functional TADs in their N-terminal region capable of interacting with the MED15 KIX domain. We further show that the ability to co-activate gene transcription by DELLA proteins involves the recruitment of Mediator complex through MED15 in different physiological processes, and propose that this mechanism evolved in a common ancestor of land plants.

## Results

### DELLA proteins recruit MED15 to promote transcription

We have previously shown the extensive conservation of a predicted nine amino acid (9aa) transcription activation domain (TAD) located within α-helix D of DELLA proteins (**Fig. 1A, *SI Appendix*, Fig. S1**, Hernández-García et al., 2019). After reanalysing the N-terminal sequence, which includes the first 120 amino acids of DELLAs, we found additional potential 9aaTADs that overlap with the canonical DELLA and LEQLE motifs. Together with that found in α-helix D, these 9aaTADs encompass ΦxxΦΦ motifs surrounded by acidic residues, where Φ are bulky hydrophobic residues (**Fig. 1A**). These motifs are essential for the function of the activation domains of eukaryotic transcription factors such as the mammalian tumour suppressor p53 (25, 26). To assess the functional significance of the predicted motifs in DELLA-mediated transcriptional activation, we used a GAL4-UAS-based yeast assay. While the GAL4 DNA binding domain (GAL4BD) fusion to a fragment of RGA containing the first 120 aa (named N2) caused activation of the reporter, mutation of the Φ residues within the predicted motifs into alanine residues abolished this activity (**Fig. 1B**), indicating that these motifs are indispensable for transcriptional activation in a heterologous system. To confirm if this holds true in plant cells, we used a plant transient expression Gal4-based dual luciferase system(27) with an enhanced 5xUAS Gal4-binding promoter driving firefly luciferase expression. With this system, GAL4BD-RGA^N2^ robustly induced luciferase expression six-fold compared to a control, while the Φ-to-A mutant (GAL4BD-RGA^N2-mΦ^) experienced a drastic reduction of 85-95% of the activity (**Fig. 1C**), suggesting that ΦxxΦΦ-type TADs are the main elements governing DELLA-mediated transcriptional activation in yeast and plants.

**Figure 1.**
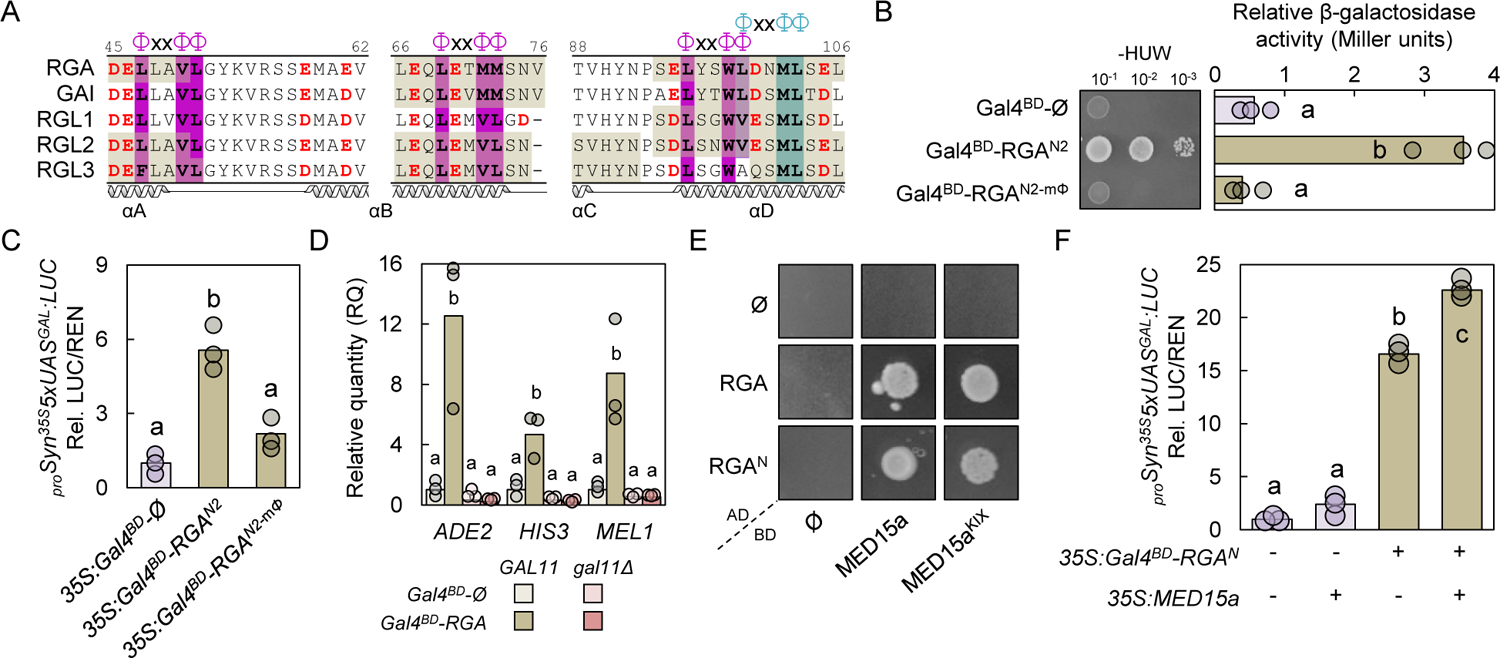
RGA and MED15 interaction enhances gene activation. **A)** Alignment of Arabidopsis DELLA selected regions (numbers above alignment refer to corresponding RGA residue) showing the presence of predicted 9aaTADs (khaki shaded residues), p53-like activation domain motif bulky residues (Φ, purple shading, or cyan shading in overlapping motif), and acidic residues (red) surrounding p53-like AD sites. ‘x’ refers to any residue. Secondary structure showing α helixes based on Murase et al. 2008 (77). B) Yeast one-hybrid transactivation assay of GAL4 DNA binding domain (Gal4^BD^) fusions of RGA N2 region (amino acids 1-120) with the Φ residues shown in A) mutated to Ala residues or not (see **Fig. S3D**). The left side shows a drop assay of a dilution series on - His medium and the right-side quantification of the LacZ reporter activation. C) Dual luciferase transactivation assay in *Nicotiana benthamiana* leaves using the *LUC* gene under the control of 5xGal operon UAS motif as the reporter, and either a Gal4^BD^ fused to the RGA N2 region with Φ residues mutated or not. D) RT-qPCR analysis of the GAL4-responsive reporter genes in the original Y2HGold yeast strain (*GAL11*) or with the endogenous *MED15/GAL11* gene deleted (*gal11Δ*) in the presence of a Gal4^BD^ alone or fused to RGA. *Saccharomyces cerevisiae ACT1* was used as reference gene. E) Yeast two-hybrid assay using MED15a or MED15a KIX domain as bait, and RGA and RGA DELLA domain as prey. F) Dual luciferase transactivation assay in *N. benthamiana* leaves using the *LUC* gene under the control of 6xGal operon UAS motif as the reporter, and a Gal4^BD^ effector fused to the RGA-N co-expressed or not with MED15a as co-effector. Bars represent the mean activity of three independent biological replicates (circles). Statistical groups in determined by Tukey’s Post-Hoc test (p<0.05) following ANOVA analysis. For RT-qPCR, groups were calculated independently per analyzed gene.

The presence of 9aaTADs has been associated to the interaction with KIX domains through these regions (26). Such domains are present in various eukaryotic transcriptional co-activators as the Mediator tail subunit MED15 and the p300/CBP family of histone acetylases (26). Given the absence of a KIX domain in the single p300/CBP homolog in yeasts (28), we focused on the potential recruitment of MED15 by DELLA proteins as a mechanism to induce transcriptional activation. Supporting this hypothesis, CRISPR-mediated knock-out of *GAL11*, the single *MED15* gene in *Saccharomyces cerevisiae*, prevented RGA-mediated transcriptional activation of the endogenous reporters in our experimental strain (**Fig. 1D**). In addition, RGA exhibited interaction by yeast two-hybrid (Y2H) assays with Arabidopsis’ main MED15 protein, MED15a, that depended on both the RGA N-terminal (RGA^N^) and KIX domain (MED15a^KIX^) (**Fig. 1E, *SI Appendix*, Fig. S2A**). In these assays, the α-helix D of RGA alone exhibited a modest, yet discernible interaction with MED15a^KIX^, pointing to the potential role of this region to mediate the interaction. We confirmed this interaction via Bimolecular Fluorescence Complementation (BiFC) assays in *Nicotiana benthamiana* (***SI Appendix*, Fig. S2B**), and extended it to additional paralogs of both proteins, namely MED15d and MED15f, alongside DELLA proteins GAI and RGL2 (***SI Appendix*, Fig. S2C-D**). We additionally observed partial loss of interaction between the KIX domain and RGA versions lacking α-helix D (*rga^ΔαD^*), yet not with *rga^Δ17^*, an RGA mutant version missing its canonical DELLA motif together with the next 12 downstream amino acids (***SI Appendix*, Fig. S3A-B**). This further points to the interaction mainly relying on the presence of the α-helix D, while also suggesting the potential presence of additional, and weaker, KIX-interacting motifs within RGA. Subsequent activation assays revealed that *rga^Δ17^* retained 78% of the non-deleted version transcriptional activation capability, the α-helix D deletion (*rga^ΔαD^*) retained around 30% of it, and the combined deletion of both regions (*rga^2Δ^*) completely abolished the ability to induce reporter activation (***SI Appendix*, Fig. S3C**). This implies that DELLA’s capability to induce transcription is encoded exclusively in the N2 region, with the α-helix D being the main - but not the only-region associated with it. Accordingly, the interaction between RGA-N2 and MED15a^KIX^ was completely impaired upon mutation of the ΦxxΦΦ motifs (***SI Appendix*, Fig. S3D**). To assess whether this interaction mediates transcriptional activation in plant cells, we introduced MED15 together with GAL4BD-RGA^N^ in our adapted *N. benthamiana* transient dual luciferase system. We observed that the addition of MED15a further increased the ability of RGA^N^ to activate transcription (**Fig. 1F**). Altogether, these findings suggest that DELLAs would recruit Mediator complexes to induce transcriptional activation, pointing to a direct role of DELLA proteins as transcriptional co-activators.

### MED15 is necessary for the activation of a subset of DELLA-dependent transcriptional responses

To assess the biological relevance of DELLA-MED15 interaction, we analysed the DELLA-dependent transcriptome on a RNAi line targeting *MED15a* (termed here *med15a^Ri^*, Kim et al., 2016). We compared the transcriptomic profile of wild-type (WT) and *med15a^Ri^* whole seedlings with high vs low DELLA levels (i.e., treated with the GA-biosynthesis inhibitor paclobutrazol [PAC] or with PAC + GA_3_, respectively, see **Materials and Methods**). We searched in the WT under high DELLA levels (PAC treatment) for genes differentially induced, which were not induced (or induced at significant lower levels) in the *med15a^Ri^* line. Then, we selected genes with at least 1 Transcript per Million Reads (TPM) in both genotypes, and that were differentially expressed between high and low DELLA levels (in PAC vs PAC+GA_3_, p-adj < 0.01). A total of 681 genes were induced at high DELLA levels in WT plants and 1737 in the *med15a^Ri^* line (**Dataset S1**). Among the genes induced in WT plants, 497 were not induced in the *MED15a* RNAi line (**Fig. 2A**), in line with a mechanism of DELLA-dependent gene induction relying on MED15 function. Up to 1548 (89%) of the genes up-regulated in *med15a^Ri^* were not induced in WT, suggesting that impaired Mediator activity increases the sensitivity towards high levels of DELLA protein. More importantly, 183 out of 184 genes upregulated by DELLAs in both genotypes were more weakly induced in the *med15a^Ri^* line when comparing their absolute expression, contrasting with the expected opposite pattern in the subset of genes up-regulated only in *med15a^Ri^* (**Fig. 2B, *SI Appendix*, Fig. S4**). We then define the list of up-regulated genes in WT but not in *med15a^Ri^* as the putative DELLA-MED15 target candidates’ list.

**Figure 2.**
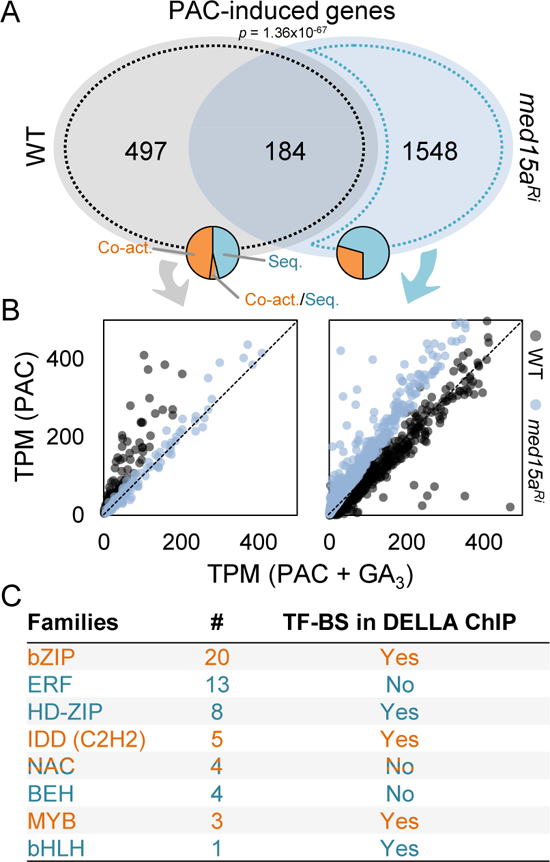
MED15 is involved in DELLA-mediated transcriptional activation. **A)** Venn’s diagram showing the overlap of curated genes (showing at least 1 TPM in WT and *med15a^Ri^*, padj <0.01) up-regulated in PAC-treated plants (compared to PAC+GA_3_-treated plants). p-value shown indicates the statistical significance of the overlap. Orange-blue circles represent the predicted DELLA mechanistic effect on EAT-Up-derived TFs found to be enriched in each subset. Orange, co-activation; blue, sequestering. B) TPM comparison of the genes up-regulated in WT (grey dot line in A), or exclusively in *med15a^Ri^* (blue dot line in A). Zoomed-out axes are shown in Fig. S4. C) TF numbers per family and the presence of TF binding sites in DELLA ChIP-seq experiments (Marín-de la Rosa et al., 2015). Colours represent the predicted DELLA mechanistic effect on that TF family. Expression analyses were carried out by whole RNA sequencing of 7-day-old seedlings treated for 3 days either with 1 μM PAC, or 1 μM PAC + 100 μM GA_3_.

DELLAs are unable to directly bind DNA, instead they are brought to chromatin through the interaction with different DNA-binding TFs (29). We looked for predicted TFs involved in the regulation of the putative DELLA-MED15 targets using the EAT-UpTFv0.1 tool to find enriched TF binding sites (BS) in their upstream regulatory sequences (30). We found 52 TFs with BS significantly enriched in the regulatory sequences of the putative DELLA-MED15 targets, all belonging to 8 TF families known to be direct interactors of DELLAs (**Fig. 2A,C, Dataset S2**). These include four out of the five TF families that have been reported to require DELLA as co-activators: INDETERMINATE DOMAIN (IDD) proteins of the C2H2 family, MYBs, NACs, and bZIPs (12, 13, 31–34). In addition, five of the eight families bind to *cis* elements enriched in DELLA-binding regions previously identified by ChIP-seq (3, 35). In contrast, an equivalent analysis with the upstream sequences of the genes upregulated in the *med15a^Ri^* line yielded enrichment for only 17 TFs belonging to 7 different TF families. Of these TFs, 12 are predicted to be regulated by DELLAs through sequestration, and 2 belong to families that have never been associated with DELLA activity (**Fig. 2A**). This analysis indicates that the DELLA-MED15 target candidates are genes whose up-regulation preferentially depends on DELLA’s recruitment to the DNA through specific DELLA-TF interactions. Moreover, this observation suggests TFs known to act with DELLAs as co-activators regulate DELLA-MED15 target genes.

### MED15 is required for DELLA regulation of biological processes as a co-activator

Among the GO terms enriched within the putative DELLA-MED15 targets, we found “response to GA stimulus” (GO:0009739), together with other GA-mediated processes as “response to oxidative stress” (GO:0006979), and “secondary metabolism” (GO:0019748), mostly due to the regulation of flavonol biosynthesis (***SI Appendix*, Fig. S5, Dataset S3**). These are processes for which DELLA has been shown to act as co-activators of IDD and MYB12 TFs, respectively (12, 13, 32). Conversely, we found “growth” (GO:0040007), or “response to cold” (GO:0009409) between the GOs enriched in the genes up-regulated only in the *med15a^Ri^* line. These terms can be associated with known GA responses, but no evidence links any of these biological processes to co-activation. This prompted us to propose that MED15 may be required to regulate a subset of previously known GA transcriptional responses.

To test this hypothesis, we examined the impact of reducing MED15 activity in contexts where DELLAs act either sequestering TFs or as co-activators. The concerted function of the PIF, BZR and ARF families of TFs in hypocotyl elongation is counteracted by direct DELLA-sequestering of all these effectors (36). We observed an equal reduction of hypocotyl elongation in the presence of PAC in WT and *med15a^Ri^* seedlings, (***SI Appendix*, Fig. S6A-B**). In line, we found no change in the expression levels on the PIF-repressed genes *PIL1* and *XTR7,* or the PIF-induced gene *PRE5* (***SI Appendix*, Fig. S6C**). Similarly, hypocotyl elongation reduction by PAC in plants lacking the Mediator tail and head subunits MED5 and MED8 (*med5ab* and *med8*) was comparable to that of WT (***SI Appendix*, Fig. S6D**). We then tested the involvement of MED15 in biological processes regulated by DELLA co-activation together with TFs. A well-documented process is DELLAs regulation of developmental programs in hypocotyls and apical meristems acting as co-activators of type-B ARRs (3). DELLA accumulation promotes cotyledon opening during skotomorphogenic development (37). In agreement with a requirement for MED15 in DELLA co-activation, we observed a defective response to PAC-induced cotyledon opening in the *med15a^Ri^* line (**Fig. 3A-B**). Interestingly, we found defective DELLA-mediated cotyledon opening in darkness also in the *med5ab* and *med8* mutants (***SI Appendix*, Fig. S7A**), pointing to a generalized function of the whole complex mediating this response. In addition, we observed that the *med15a^Ri^* line was impaired in DELLA-mediated reduction of root apical meristem size (***SI Appendix*, Fig. S7B**), another process regulated by co-activation of DELLAs with type-B ARRs (3). Flavonol production has been reported to be promoted by DELLAs through concerted co-activation with MYB12 (13). In line with our hypothesis, PAC-dependent flavonoid accumulation in roots of skotomorphogenic plants was evident in WT root tips but severely impaired in *med15a^Ri^* plants, which was in line with the inability to fully activate transcription of the *CHS* and *FLS1* genes (**Fig. 3C, *SI Appendix*, Fig. S8A**). A third case of TFs interacting with DELLA proteins to regulate the GA negative feedback is the IDD family, which directly up-regulates GA biosynthetic genes in response to increased DELLA levels (32). Accordingly, PAC-dependent induction of *GA20ox2* and *GA3ox1* expression was significantly reduced in the *med15a^Ri^* line compared to WT plants (**Fig. 3D**). A direct target of IDD1 and other IDD proteins involving DELLAs co-activation is *SCL3* (12), for which we observed a robust induction after DELLA accumulation which was severely impaired under MED15 reduced function (**Fig. 3D**). Altogether, these results indicate that MED15, and likely the whole Mediator complex, is involved in the gene regulation of developmental processes mediated by TF-DELLA transcriptional co-activation, but not for processes regulated by TF-DELLA sequestering mechanisms.

**Figure 3.**
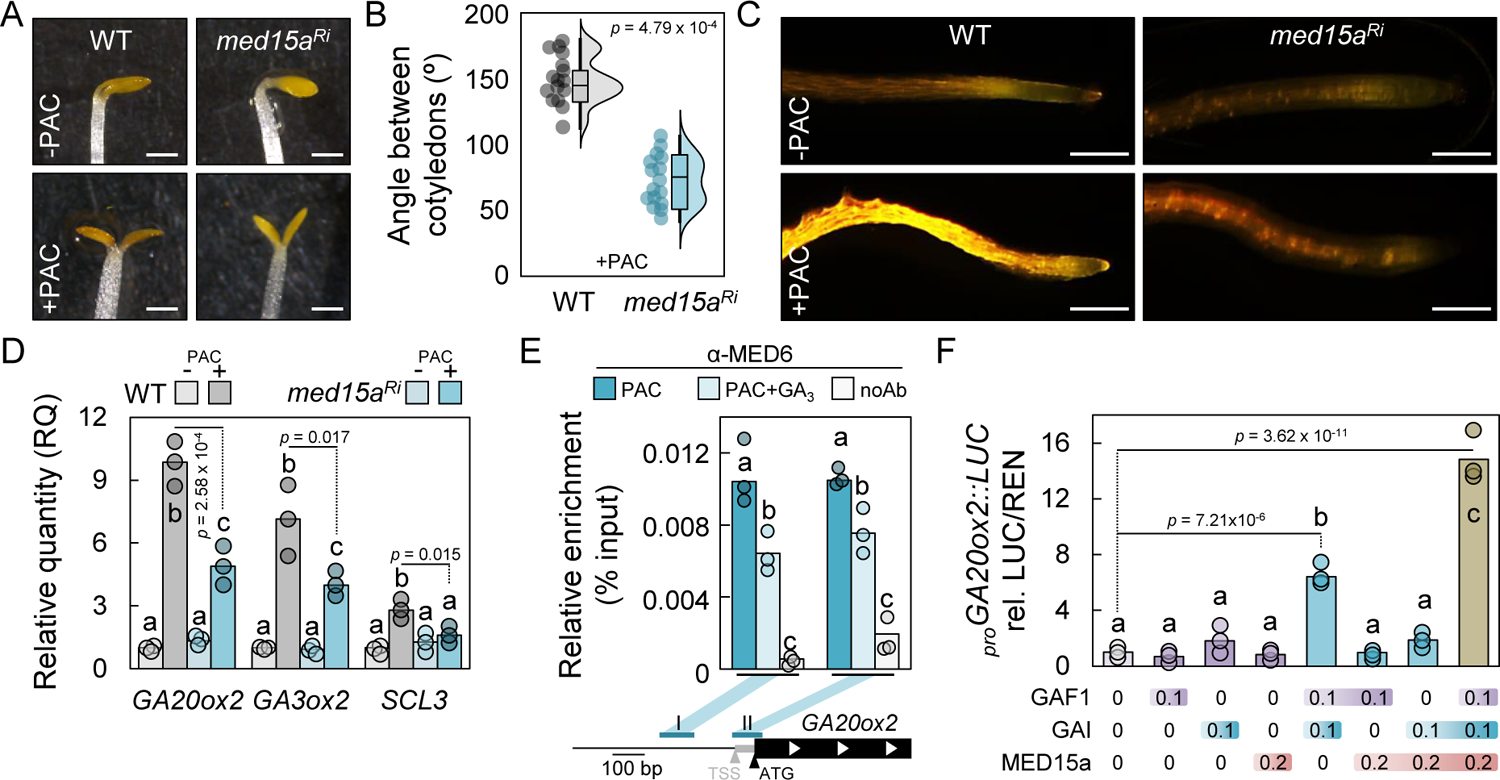
MED15 is needed for DELLA-dependent co-activation responses. **A)** 5-day-old WT and *med15a^Ri^* seedlings grown in darkness with or without 1 μM PAC. Scale bar, 1 mm. B) Rainbow plot of angle between cotyledons in 5-day-old WT and *med15a^Ri^* seedlings grown in darkness with or without 1 μM PAC. Measurements in mock plants are not shown (mean = 0; SD = 0). C) DPBA staining of 8-day-old WT and *med15a^Ri^* seedlings root tips grown in long day conditions (16L:8D), and treated for four days with or without 1 μM PAC. Scale bar, 100 μm. D) RT-qPCR analysis of GA and IDD-responsive genes in 5 days-old WT and *med15a^Ri^* seedlings grown in darkness with or without 1 μM PAC using *PDF2.1* as reference gene. E) ChIP-qPCR analysis of MED6 occupancy at the indicated locations of *GA20ox2* promoter in 7-day-old seedlings treated with 1 μM PAC, and with or without 100 μM GA_3_ for 3 days. Relative enrichment is given as a percentage of IP/Input. F) Dual luciferase transactivation assay in *N. benthamiana* leaves using the *LUC* gene under the control of *GA20ox2* promoter as the reporter, and HA-GAF1, YFP-GAI, and cMyc-MED15a co-expressed as effectors. The effector level is indicated below each bar as the agroinfiltrated OD_600_. B) shows experimental data of one representative experiment of three with at least 15 plants per genotype and treatment; statistical difference represented as Student’s t-test *p*-value. D-F) bars represent biological replicate means and means of technical triplicates are depicted as circles. Statistical groups in D-F) were determined by Tukey’s Post-Hoc test (p<0.05) following ANOVA analysis. For qPCR, ANOVA analyses were performed independently per gene/genomic region.

If Mediator is recruited to chromatin contexts by DELLA to induce transcription, promoter regions in DELLA-induced genes targeted by TF-DELLA complexes should also be targeted by Mediator. The *GA20ox2* and *SCL3* promoters are bound by both IDDs and DELLAs to induce their transcription (12, 32, 35). We thus performed chromatin-immunoprecipitation assays with available anti-MED6 antibodies (38) under increased or depleted DELLA levels (*i.e.* PAC or PAC+GA_3_-treated WT plants, respectively). By quantitative PCR, we found that MED6 was significantly enriched under increased DELLA levels in the two analysed *GA20ox2* promoter regions, and in the proximal region of *SCL3* promoter (II), coinciding with the regions bound by IDDs and DELLAs(32, 35), and supporting the role of DELLAs in recruiting Mediator to these loci (**Fig. 3E, *SI Appendix*, Fig. S8B**). Furthermore, we tested the transcriptional output of the *in vivo* reconstructed GAF1-DELLA-MED15 module in a dual luciferase assay. When co-expressed, GAF1 and either GAI and RGA could induce *GA20ox2* promoter activity as previously shown (32), and the addition of MED15a was sufficient to enhance luciferase activity in a DELLA and GAF1-dependent way (**Fig. 3F, *SI Appendix*, Fig. S8C**). Altogether, these results show that IDD-DELLA activation of gene expression requires MED15 direct recruitment to specific target loci to regulate a subset of transcriptional responses.

### DELLA recruitment of MED15 is a conserved mechanism in land plants

The conservation of the DELLA activation domain motifs (***SI Appendix*, Fig. S1**) suggests that DELLA co-activation through Mediator recruitment may occur in all land plants. We therefore decided to test the function as co-activator of the single DELLA protein in the liverwort *Marchantia polymorpha* (MpDELLA). It has been previously shown that MpDELLA can act through TF-sequestration, and that its N-terminal domain has intrinsic transactivation function in yeasts (8, 15). We identified a single full-length *MED15* gene in *M. polymorpha*, *Mp8g02180* (hereafter referred to as Mp*MED15*, ***SI Appendix,* Fig. S9, Materials & Methods**), whose KIX domain interacted with MpDELLA in Y2H assays, (**Fig. 4A, *SI Appendix*, Fig. S10A**). Our BiFC analysis indicated that MpDELLA-MpMED15 interaction occurs in the nucleus, and confirmed the requirement of MpDELLA N-end for the interaction (***SI Appendix*, Fig. S10B**). We next tested if the MpDELLA-MpMED15 interaction resulted in the activation of transcription *in vivo*. First, we confirmed that MpDELLA N-terminal (MpDELLA^N^) domain alone is able to activate gene expression in plant cells using our Gal4-based dual luciferase assay (**Fig. 4B, *SI Appendix*, Fig. S10C**). The addition of MpMED15 consistently enhanced the ability of MpDELLA^N^ to induce LUC activity (**Fig. 4B**), echoing the effect of Arabidopsis proteins, and supporting a conserved mechanism of DELLA recruiting MED15 to promote transcription in land plants.

**Figure 4.**
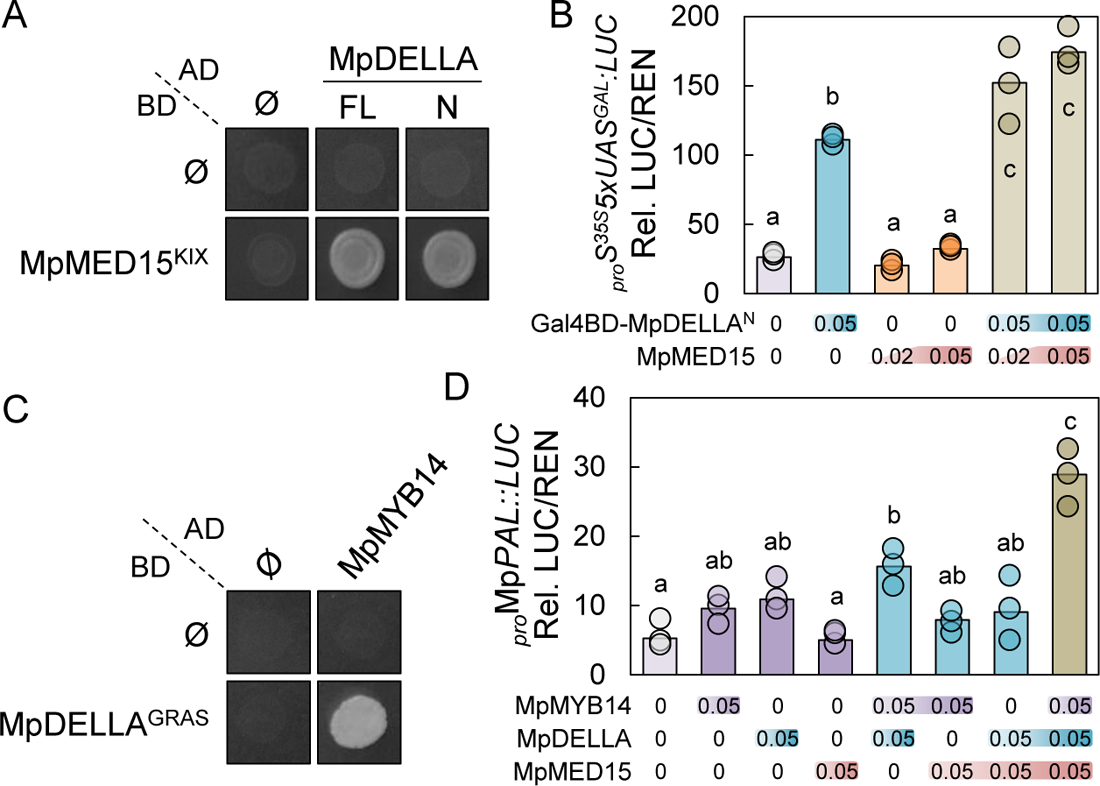
Marchantia polymorpha MpDELLA acts as a co-activator by recruiting MED15. A) Yeast two-hybrid assay using MpMED15 KIX domain as bait, and MpDELLA full length (FL) or N-terminal domain (N) as prey. B) Dual luciferase transactivation assay in *Nicotiana benthamiana* leaves using the *LUC* gene under the control of the *Gal* operon UAS promoter as the reporter, and a GAL4 DNA binding domain (DBD) fused to the MpDELLA N-terminal domain, and MpMED15 co-expressed as effectors. C) Yeast two-hybrid assay using MpDELLA GRAS domain as bait, and MpMYB14 as prey. D) Dual luciferase transactivation assay in *N. benthamiana* leaves using the *LUC* gene under the control of Mp*PAL* (*Mp1g05190*) promoter as the reporter, and HA-FLAG-MpMYB14, YFP-MpDELLA, and cMyc-MpMED15 co-expressed as effectors. Effector level in C and E is indicated below each bar as the agroinfiltrated OD_600_. Bars represent medians of 3 biological replicates. Biological replicate means of technical triplicates are depicted as circles. Statistical groups were determined by Tukey’s Post-Hoc test (p<0.05) following ANOVA analysis.

To confirm that MpDELLA-MpMED15 interaction could be relevant in the context of a *M. polymorpha* promoter. We focused on processes regulated by DELLAs and TFs by co-activation that could be conserved in *M. polymorpha*, such as flavonoid biosynthesis (8, 39, 40). MpDELLA has been shown to interact with orthologs of MpMYB14 (9), and both share common activated transcriptional targets as Mp*PAL* (*Mp1g05190*). This led us to hypothesize that MpMYB14 and MpDELLA would work together to engage MpMED15 and promote gene transcription. First, we confirmed the interaction between MpDELLA and MpMYB14 by Y2H and BiFC (**Fig. 4C, *SI Appendix*, Fig. S11**). This interaction requires the GRAS domain of MpDELLA and occurs in plant nuclei. In agreement with previous results, MpMYB14 could consistently activate Mp*PAL* transcription using a *_pro_*Mp*PAL::LUC* reporter in a dual luciferase assay (***SI Appendix*, Fig. S12A**). Supporting our hypothesis, MpDELLA significantly promoted MpMYB14 ability to activate Mp*PAL* transcription (***SI Appendix*, Fig. S12B,C**). This suggests that MpDELLA cooperates with MpMYB14 as a co-activator of its target genes, at least in the case of Mp*PAL*, indicating that MpDELLA functions as a TF co-activator also in *M. polymorpha*. The addition of MpMED15 to the system further increased luciferase activity in a MpDELLA and MpMYB14-dependent way (**Fig. 4D**). Altogether, these results support the idea that the recruitment of Mediator by DELLA is a conserved mechanism of transcriptional co-activation in land plants.

## Discussion

The findings described here link DELLA’s reported function as transcriptional coactivators to the recruitment of the Mediator complex through physical interaction between conserved TADs in the N-terminal end of DELLA and the KIX domain of MED15. In support of the relevance of this interaction, our molecular analysis shows on the one hand that Mediator’s recruitment to certain DELLA target genes requires the concurrence of DELLA itself, and functional studies on the other hand indicate that MED15 is required for an optimal response to DELLA accumulation. The N-terminal domain of angiosperm DELLAs contains up to four TADs, but only the two present in the α-helix D seem critical for the interaction with MED15, coinciding with the deeply conserved TADs in all land plant DELLAs. A third TAD overlapping with the DELLA motif contributes to a lesser stent, and evolved later during angiosperm evolution. The presence of multiple functional TADs has been shown to be important for multiple eukaryotic transcriptional activators (21, 41, 42).

Transcriptional activation has also been described to occur in eukaryotes through mechanisms not involving Mediator. For example, metazoan TFs also activate transcription via the interaction between their TADs and the p300/CBP histone acetyltransferase (HAT)-type KIX domains (26). The ability of DELLA to activate transcription in yeast –which lacks KIX-containing HATs– favors MED15 as the relevant partner for co-activation. However, plant genomes encode KIX-containing HATs (43), and some TFs as SREBP1a have been shown to alternatively recruit CBP or MED15 in different situations (44). Thus, DELLAs might recruit HATs to activate transcription in plant cells through a Mediator-independent mechanism. Our transcriptomic analysis unexpectedly showed that reducing MED15 activity causes a large set of genes to be more responsive to DELLA than in wild-type plants. One possible explanation would be that an impairment in Mediator activity would be compensated by reinforcing the interaction between DELLA and putative Mediator-independent transcriptional regulators. However, HAC1 and HAC5, the KIX-containing p300/CBP orthologs in Arabidopsis, have been proposed to be part of the plant Mediator complex (45), suggesting that a putative DELLA-HAC interaction in plants would ultimately lead to the recruitment of Mediator.

At least two pieces of evidence suggest that the function of DELLA as a transcriptional coactivator is an ancestral feature: (i) the robust conservation of TAD sequences in the N-terminal region of DELLAs from all extant land plants (15); and (ii) the functional conservation of MpDELLA and MpMED15 interaction shown here (**Fig. 4A, *SI Appendix*, Fig. S12**). Thus, it is likely that both mechanisms of DELLA-dependent transcriptional regulation –TF sequestration and co-activation– are intimately linked to the protein structure and have coexisted in DELLAs since their origin in a last common ancestor of land plants. Of note, DELLAs might also act in certain contexts as active repressors by a yet unknown mechanism (46), and future research will be needed to clarify how and why DELLAs vary their transcriptional activity in a context-dependent manner. Whether one mechanism is more relevant than the others in specific biological processes is hard to discern at this point.

An interesting observation is that the region containing the TADs overlaps with the GID1 receptor binding region. GID1 binds the DELLA domain in the presence of GAs, triggering DELLA ubiquitylation and subsequent de-stabilization; thus, this region can be considered the GA-dependent degron (47). The overlap of degrons and TADs in the same region has been documented in several yeast and mammalian TFs, such as GCN4, c-Myc, or p53 (48). It has been proposed that this coincidence allows quick destabilization of transcriptional activators, an advantage for the fast reprogramming of transcriptional responses by a sequential mechanism of fast and transient blockage and subsequent, slower and irreversible degradation. In agreement with this, GA-dependent recognition of DELLA by the GID1 receptor has been shown to block transcriptional activation in yeast without involving DELLA degradation (14). Assuming that the last common ancestor of bryophytes and tracheophytes did not possess GID1 GA receptors (15), it is tempting to speculate that the incorporation of DELLA to GA signaling as we know it in extant tracheophytes occurred in two steps: the establishment of GA-GID1-DELLA interaction to block the activity of TADs in DELLA’s N-terminal region by competition with MED15, and then the recruitment of the SCF^SLY1^ complex for DELLA polyubiquitylation. However, we lack evidence that supports the existence of a plant where DELLA is only regulated by GID1-mediated impairment of its transactivation activity, or that shows that this mechanism is relevant in specific biological contexts.

## Materials and Methods

### Transcriptional activation domain prediction

Transcriptional activation domain prediction was performed on previously reported DELLA sequences (15) with the 9aaTAD prediction tool (http://www.med.muni.cz/9aaTAD/index.php) using the “less stringent” pattern (49), and automatically searching for p53-like TADs with the ΦxxΦΦ motif sequence, were Φ are bulky hydrophobic amino acids and x any amino acid, with at least one acidic residue in close vicinity (±3 residues). The protein sequences were aligned with MAFFT 7.0, using the L-INS-I method (50), followed by manual curation.

### Yeast transactivation assays

Gal4-DNA Binding Domain (BD) fusions were made by transferring the indicated DELLA versions into the pGBKT7-GW vector (51). New entry vectors were obtained by transferring PCR-amplified, or gBlock (I.D.T.) synthesized, CDSs into pDONR207 via BP Clonase II (Invitrogen). Deletions were performed in the entry vectors by PCR and blunt-end re-ligation. Final constructs were made by recombining entry clones with Gateway destination vectors via LR Clonase II (Invitrogen), and transformed into the *S. cerevisiae* strain Y2HGold and selected in SD medium lacking Trp. Drop tests were performed in solid SD medium without Trp and His, and increasing 3-aminotriazol concentrations as indicated. For quantitative assays, the GBKT7 containing haploid Y2HGold strains were mated with a *S. cerevisiae* Y187 strain bearing a disarmed plasmid conferring Leu selection, and selected in SD medium lacking Leu and Trp. Diploid strains were grown in liquid SD and β-galactosidase activity measurements were performed as previously described (52) using ortho-nitrophenyl-β-galactoside (ONPG) as a substrate.

### Yeast-two hybrid assays

For bait Gal4-DNA Binding Domain (BD) fusions, MED15-related sequences (full length, KIX domain and C-terminal domain), and a truncated version of MpDELLA (MpDELLA^Δ2^) based on previously described DELLA deletions were introduced in the pGBKT7-GW vector (51). DELLA full-length, N-terminal and C-terminal sequences from Arabidopsis and Marchantia, and MpMYB14 were fused to the Gal4-Activation Domain (AD) in pGADT7-GW (51). New entry vectors were obtained by transferring PCR-amplified CDSs to pDONR207 via BP Clonase II (Invitrogen). Final constructs were made by recombining entry clones with Gateway destination vectors via LR Clonase II (Invitrogen). Direct interaction assays in yeast were performed following Clontech’s small-scale yeast transformation procedure. Strain Y187 was transformed with pGADT7-derived expression vectors, while strain Y2HGold was transformed with pGBKT7 vectors, and selected in SD medium without Leu or Trp, respectively. Subsequently, diploid cells were obtained by mating and selection in SD medium lacking both Leu and Trp. Drop interaction tests were done in SD medium lacking Leu, Trp and His, in the presence of different concentrations of 3-aminotriazol (3-AT) (Sigma-Aldrich). Quantitative interaction assays were performed in liquid medium by quantifying β-galactosidase activity as previously described (52).

### Bimolecular fluorescent complementation (BiFC) assays

For BiFC, DELLA-related entries, and MED15/MpMYB14 entries were recombined with pMDC43-YFN and pMDC43-YFC (53), respectively. *Agrobacterium tumefaciens* GV3101 containing binary plasmids were used to infiltrate 4-week-old *Nicotiana benthamiana* leaves. Three days after infiltration, leaves were analysed with a Zeiss LSM 780 confocal microscope. Reconstituted YFP signal was detected with emission filters set to 503-517 nm. Nuclei presence in abaxial epidermal cells was verified by transmitted light.

### Dual luciferase transactivation assays

For Gal4-based assays, a reporter construct containing 5xGal4 UAS flanked by an upstream 35S promoter enhancer and a downstream 35S minimal promoter was amplified from the pMpGWB101-5xUAS:LUC plasmid (54) using sequence-specific primers and introduced upstream of the firefly luciferase gene (LUC) in the pGreenII 0800-LUC plasmid (55) by a NEBuilder HiFi DNA Assembly reaction (New England Biolabs) to create the *_pro_Syn^35S^5xUAS^GAL^:LUC* plasmid also containing a *_pro_35S:REN* cassette. For Gal4 DNA binding domain fusion to RGA^N2^ and the TAD mutant version (Fig. 1C), these parts were transferred into the pMpGWB102-GAL4DBD plasmid (54) via LR Clonase II (Invitrogen) from the entry vectors. The Gal4^BD^-RGA^N^ effector plasmid has been previously described (15). AtMED15a/MpMED15, full-length RGA/GAI/MpDELLA, and GAF1/MpMYB14 constructs were introduced into pEarleyGate203, pEarleyGate104 and pEarleyGate201 destination vectors (56) via LR Clonase II (Invitrogen) from their entry vectors, respectively. The Gal4^BD^-MpDELLA^N^ effector vector was obtained by amplifying the GAL4 BD-MpDELLA N-end fusion previously generated in the pGBKT7 vector as a unique PCR product with proper restriction site overhangs and ligated into pFGC5941 (http://www.ChromDB.org) between the XhoI and SpeI sites. The plasmid containing the *_pro_GA20ox2::LUC* reporter was generated amplifying a previously described fragment including 1003 bp upstream and 77 bp downstream of the TSS of the Arabidopsis *GA20ox2* gene (AT5G51810) with sequence-specific primers, and cloned upstream of the firefly luciferase gene (LUC) between the SalI and BspHI sites in pGreenII 0800-LUC. The plasmid containing the *_pro_*Mp*PAL::LUC* reporter was generated amplifying a 2296 bp upstream region of the Mp*PAL* gene *Mp1g05190* from *M. polymorpha* Tak-1 genomic DNA with sequence-specific primers, and cloned upstream of the firefly luciferase gene (LUC) between SpeI and NcoI in pGreenII 0800-LUC. Transient expression in *Nicotiana benthamiana* leaves was carried out as previously reported (3). Firefly and the control *Renilla* luciferase activities were assayed in extracts from 1-cm in diameter leaf discs, using the Dual-Glo Luciferase Assay System (Promega), and quantified in a BioTek Synergy H1 Multimode Reader (Agilent, Fig. 1C), or in a GloMax 96 Microplate Luminometer (Promega, other figures). Statistical differences were analysed by one-way ANOVA followed by Tukey’s HSD post hoc test. Letters denote differences between groups with p<0.01, unless specified.

### Generation of the yeast Y2HGold *gal11Δ* mutant

The *S. cerevisiae* Y2HGold *GAL11* knockout mutant was created from the Y2HGold strain by the iterative marker-less CRISPR-Cas9 method as described previously (57, 58). The design of guide RNAs (gRNAs) for *GAL11* was carried out using the CRISPR tool within Benchling. The gRNA plasmids, gdHP013 and gdHP014 were generated by phosphorylating and annealing the primers oHP425 with oHP426 and oHP427 with oHP428, respectively. Then, a BsmBI Golden Gate reaction was performed to assemble them into the SpCas9 gRNA gap repair vector pWS2069. The donor DNA was generated by PCR-amplifying 500 bp both upstream and downstream of *GAL11* using primers oHP429-434, followed by a BsmBI Golden Gate reaction with the entry vector pYTK001. The landing pad AATGTTTCTTGTCCAAGCGGCGG was used as the unique barcode in the donor DNA. Subsequently, a BpiI-digested SpCas9 plasmid (pWS2082) with Leu selection was transformed into the Y2HGold strain along with the BpiI-digested gRNA plasmids gdHP013, gdHP014, and the donor DNA. Colony PCR was used to verify the *GAL11* knockout mutation using primers oHP435, oHP255 and oHP465, followed by the loss of Leu selection marker using YPD media.

### Plant material and growth conditions

The transgenic *med15a^Ri^* line, and the *med5ab (med5a-2/med5b-2)* and *med8-2* mutants, all in the Col-0 background, have been described before (22, 38). All seeds were surface sterilized and sown on half-strength MS w vitamins (Duchefa) plates containing 0.8% agar pH 5.7. Seedlings were grown at a constant temperature of 22 °C under long-day conditions (16 h light 60-70 μmol m^−2^s^−1^:8 h darkness) unless specified. For hypocotyl growth and cotyledon opening assessment during skotomorphogenic development, seedlings were germinated in light for 6-8 hours and transferred to darkness with or without 1 µM PAC for 5 days before evaluation. For RAM growth analysis, seedlings were grown for 5 days in vertical plates under standard conditions, and transferred to identical plates with or without 10 µM PAC for 16 hours before confocal analysis. Statistical analyses of biological samples were performed by t-test analyses between two groups, or one-way ANOVA followed by Tukey’s HSD post hoc test in multiple group comparisons. Letters denote differences between groups after ANOVA analyses (p<0.01).

### RNA isolation, cDNA synthesis, and qRT-PCR analysis

Total RNA from *S. cerevisiae* was isolated from liquid YPD cultures 8 hours after inoculation from single colonies using the formamide-EDTA method (59) followed by RNAzol^®^ RT (Sigma-Aldrich) extraction. Total RNA from *A. thaliana* 5-days-old seedlings was isolated using NucleoSpin^TM^ RNA Plant Kit (Macherey-Nagel) according to the manufacturer’s instructions. In all cases, up to 2 µg of purified RNA were treated with rDNase I (Ambion) to eliminate gDNA. cDNA was prepared from 1 μg of total RNA with NZY First-Strand cDNA Synthesis Kit (NZYTech). Quantitative real-time PCR was performed in a 7500 Fast Real-Time PCR System (Applied Biosystems) with SYBR premix ExTaq (Tli RNaseH Plus) Rox Plus (Takara Bio Inc). All relative expression levels were calculated following Hellemans et al. (2008), and *ACT1* used as reference gene (61) for *S. cerevisiae*, and *PDF2.1* (AT1G13320) for Arabidopsis (62).

### RNA sequencing and data analysis

For whole RNA sequencing, seedlings were grown under long-day conditions as described for 4 days and then transferred to either 1 µM PAC or 1 µM PAC + 100 µM GA_3_ for 3 days prior to RNA isolation. Total RNA was isolated as described above and sent to BGI Europe for quality control, library construction and sequencing on a DNBSEQ platform (BGI). The obtained 100 bp paired-end reads were analyzed with FastQC (v 0.11.9) using parameters by default to assess quality, and adaptor sequences removed with Cutadapt (with parameters ‘--minimum-length=20--max-n=0.1--quality-cutoff=30,30’) (63), and then mapped to the TAIR10 *A. thaliana* reference genome with HISAT2 (64). htseq-count was used for read count (parameters: ‘--format=bam--order=name--stranded=no’) (65), and TPMs calculated as a proxy to absolute gene expression levels. Genes with 1 TPM or more in all three replicates of a sample were considered expressed and included in the analysis of differential expression with DESeq2 v1.24.0 (66), Only differentially expressed genes (DEGs) whose adjusted p-values (p-adjust) were under 0.01 were used for metaanalyses. DEGs between PAC-treated and PAC+GA-treated seedlings can be found in **Dataset S1**. Enriched biological process terms were calculated using the GO Enrichment tool available at AgriGO using the Fisher’s test and the Yekutieli adjustment method (**Dataset S3**). For the graphic display, only Biological Process terms with p-adj values lower than 0.01 were considered and merged manually to discard duplicity in highly related terms. R code for ggplot2-based plotting can be found in the Mendeley Data link (doi: 10.17632/j474wymh93.1).

### TF binding-site and TF enrichment prediction

TF binding-site enrichment among selected genes was calculated using the <u>EAT-UpTFv0.1</u> (30). The hypergeometric statistical model with a Bonferroni post-hoc test (alpha level 0.01) was applied. The results for each of the groupings can be found in **Dataset S2**.

### Microscopy & histochemical analysis

Diphenylborinic acid 2-aminoethyl ester (DPBA) staining was used to visualize flavonoids as previously described (67). Whole seedlings were stained for 15 minutes at 0.25% (w/v) DPBA and 0.1% (v/v) Triton X-100. Epifluorescence microscopy of stained flavonoids in roots was performed on a Leica DMS1000 dissecting microscope using a GFP filter for the detection of DPBA fluorescence. Root meristems were imaged with an LSM 5 Pascal Zeiss Confocal microscope with a water-immersion objective lens (C–Apochromat 40X/1.2; Zeiss) using propidium iodide to stain cell walls. Meristem size was measured as the number of cortex cells.

### Chromatin immunoprecipitation (ChIP) and qPCR

ChIP assays were performed as previously described (68), except that samples were snap frozen in liquid nitrogen without cross-linking. Cross-linking was performed *in vitro* by adding 1% formaldehyde to the nuclear isolation buffer and incubating for 10 minutes at room temperature. Cross-linking was quenched by adding glycine to a final concentration of 125 mM, pH 8.0 before proceeding to Miracloth filtration and nuclei centrifugation. 2 g of fresh weight material were harvested per genotype in three independent biological replicates, and 2 µg of anti-MED6 (AS14-2802, Agrisera) antibody were used for immunoprecipitation. Immunoprecipitated DNA was analyzed by quantitative real-time PCR in a QuantStudio 3 system (Applied Biosystems) with SYBR premix ExTaq (Tli RNaseH Plus) Rox Plus (Takara Bio Inc). Enrichment in each sample was calculated relative to the input and expressed in percentage.

### Sequence identification and phylogenetic analysis

To identify MED15 sequences, we first built a custom BLAST database web interface based on Sequenceserver (69) with different available plant proteomes annotations: Arabidopsis thaliana Araport11 (70) was downloaded from https://www.arabidopsis.org/, *Marchantia polymorpha* v5.1 (71) was downloaded from https://marchantia.info/, *Anthoceros agrestis* Bonn (72) was downloaded from https://www.hornworts.uzh.ch/en.html, tomato iTAG4.0 (73) was downloaded from https://solgenomics.net/, *Oryza sativa* v7, *Sorghum bicolor* v3.1.1, *Physcomitrella patens* v3.3, *Selaginella moellendorffii* v1.0, *Micromonas* sp. RCC299 v3.0, and *Chlamydomonas reindhardtii* v5.5 were downloaded from https://phytozome-next.jgi.doe.gov/ (74). The in-house database was questioned by BlastP with the protein sequence of MED15a (AT1G15780) as query using an E-value cutoff of 0.1 or 10 in the case of *Micromonas* and *Chlamydomonas*. The obtained sequences were checked for the presence of KIX domains using Pfam (http://pfam.sanger.ac.uk/search). We included in our sequence list the Arabidopsis p300/CBP protein (AT1G16705) to use as an outgroup, and the identified MED15 subunits from the yeast *S. cerevisiae* (*GAL11*, YOL051W), and the amoeba *Dictyostelium discoideum* (DDB_G0293914). MED15 sequences were aligned with MAFFT 7.0, using the L-INS-I method (50), followed by manual curation. For phylogenetic reconstruction, KIX domains were used, and ambiguously aligned regions manually trimmed (see **Dataset S4**). ProtTest v3.4.2 (75) was used on final multiple sequence alignment to select the best-fit model of amino acid replacement using the AIC model for ranking. Maximum likelihood tree was produced with PhyML v3.1 (76), using the best-scored model of amino acid substitution. Statistical significance was evaluated by bootstrap analysis of 1000 replicates. Phylogenetic tree graphical representation was initially generated using FigTree (version 1.4.3) software (http://tree.bio.ed.ac.uk/software/figtree/), and final cartoon was edited manually.

### Plasmid, oligonucleotide, and other sequences

Plasmid list, primer sequences, and KIX domain aligned sequences used in this study are listed in **Dataset S4**.

## Acknowledgments

We thank Nam-Hai Chua for sharing *med15a^Ri^* line seeds, Heather Knight for sharing the *med5ab* and *med8* mutant seeds, Philip Carella and Sebastian Schornack for the pENTR-FLAG-MpMYB14 plasmid, and Takayuki Kohchi for the pMpGWB101-6xUAS:LUC and pMpGWB102-GAL4DBD plasmids. We thank the members of the Plant Signalling and Plasticity lab for their feedback on the manuscript. The research leading to these results has been carried out thanks to grants PID2019-110717GB and PID2022-141770NB (to M.A.B.) funded by the Spanish MICIN/AEI/10.13039/501100011033/ and by “ERDF, A way of making Europe”, and from European Research Council (ERC), grant DEUSBIO – 949080 (to R.L-A.).

