## Supplemental figures for "DELLA Proteins Recruit the Mediator Complex Subunit MED15 to Co-activate Transcription in Land Plants"

**This PDF file includes:**

Figures S1 to S12
Legends for Datasets S1 to S4

**Other supporting materials for this manuscript include the following:**

Datasets S1 to S4

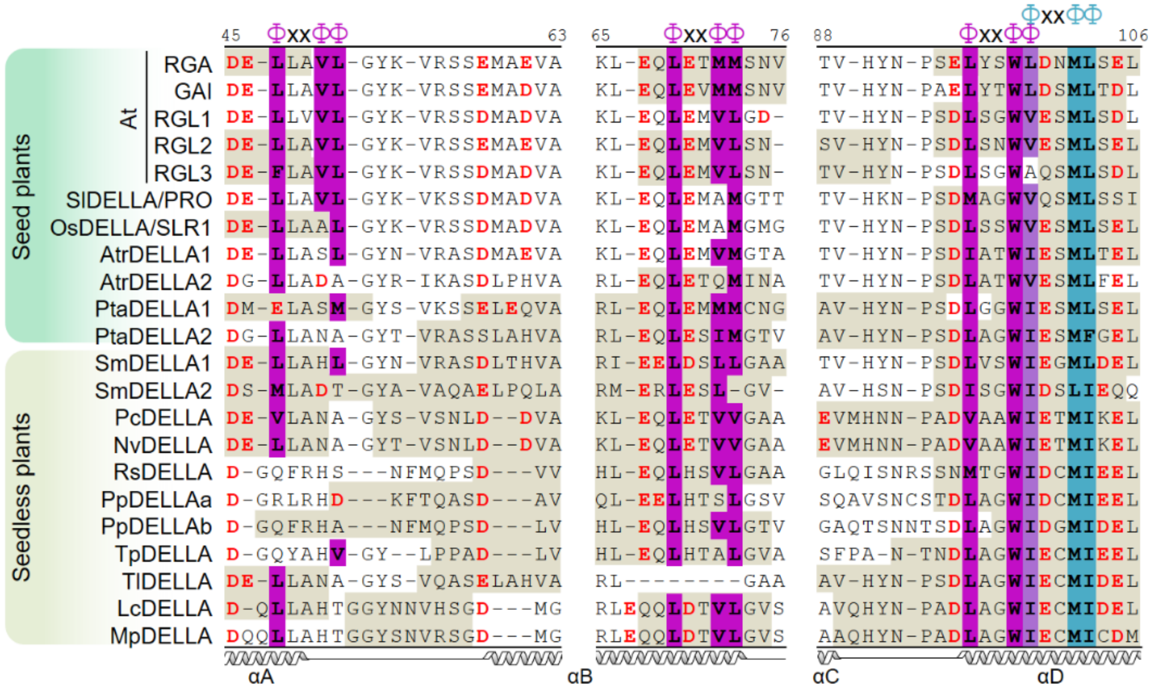


**Fig. S1. Conservation of predicted activation domains in the N-terminal motifs of DELLA proteins in multiple species**. Alignment of selected regions (number above alignment refers to corresponding RGA residue) showing the presence of predicted 9aaTADs (khaki shaded residues), p53-like activation domain motif bulky residues (Φ, purple shading, or cyan shading in overlapping motif), and acidic residues (red) surrounding p53-like AD sites. x refers to any residue. Secondary structure showing α helixes based on Murase et al. 2008. *At*, *Arabidopsis thaliana*; *Sl, Solanum lycopersicum; Os, Oryza sativa; Atr, Amborella trichopoda; Pta, Pinus taeda; Sm, Selaginella moellendorffii; Pc, Phaeoceros carolinianus; Nv, Nothoceros vincentianus; Rs, Rhyncostegium serralatum; Pp, Physcomitrium patens; Tp, Tetraphis pellucida; Tl, Takakia lepidozioides; Lc, Lunularia cruciata; Mp, Marchantia polymorpha*.

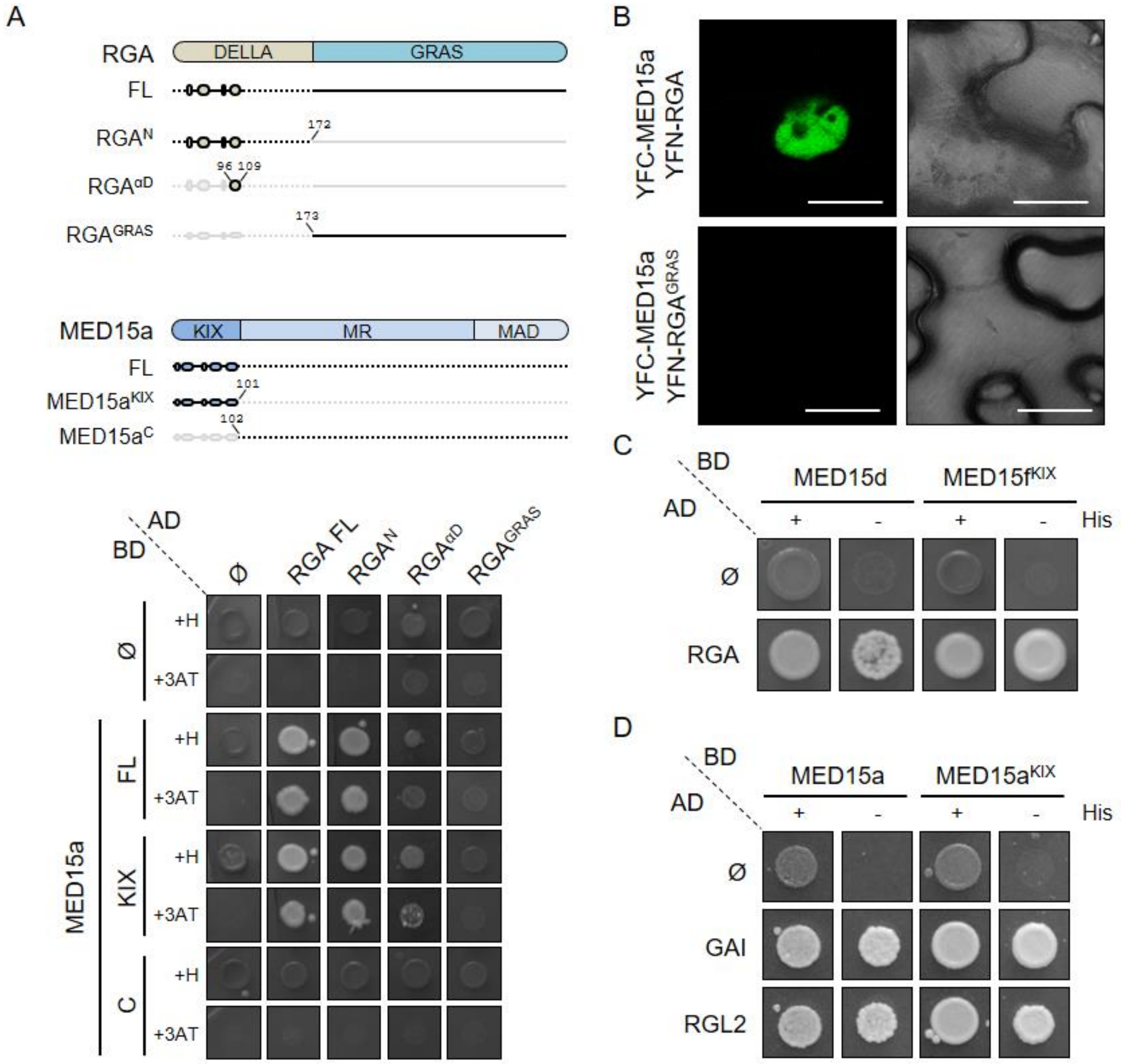


**Fig. S2. DELLAs and MED15 interact through N-terminal regions**. A) Upper panel, Illustration of RGA and MED15a deletions used. MR, Middle Region; MAD, Mediator Activation/Association Domain. Ovals represent α helixes based on known structures. Note that RGA and MED15a are not at same amino acid scale. Lower panel, yeast two-hybrid drop assay of the deletions. B) Bimolecular fluorescence complementation assay of MED15a and RGA, using a GRAS-only version as negative control. Scale bar, 10 µm. C-D) Yeast two-hybrid assays using C) MED15d and MED15f double KIX domain as baits ($Gal4^{BD}$), and RGA full length as prey ($Gal4^{AD}$), and D) MED15a as bait, and GAI and RGL2 full length versions as preys. MED15d protein is composed of two tandem KIX domains, while MED15f is a highly divergent full length MED15 subunit, encoded by a low expression gene, probably under a pseudogenization process, similarly to its closest paralog, the pseudogene *MED15e* (see Fig. S11). GAI and RGL2 are each a close and a distant paralog of RGA, respectively.

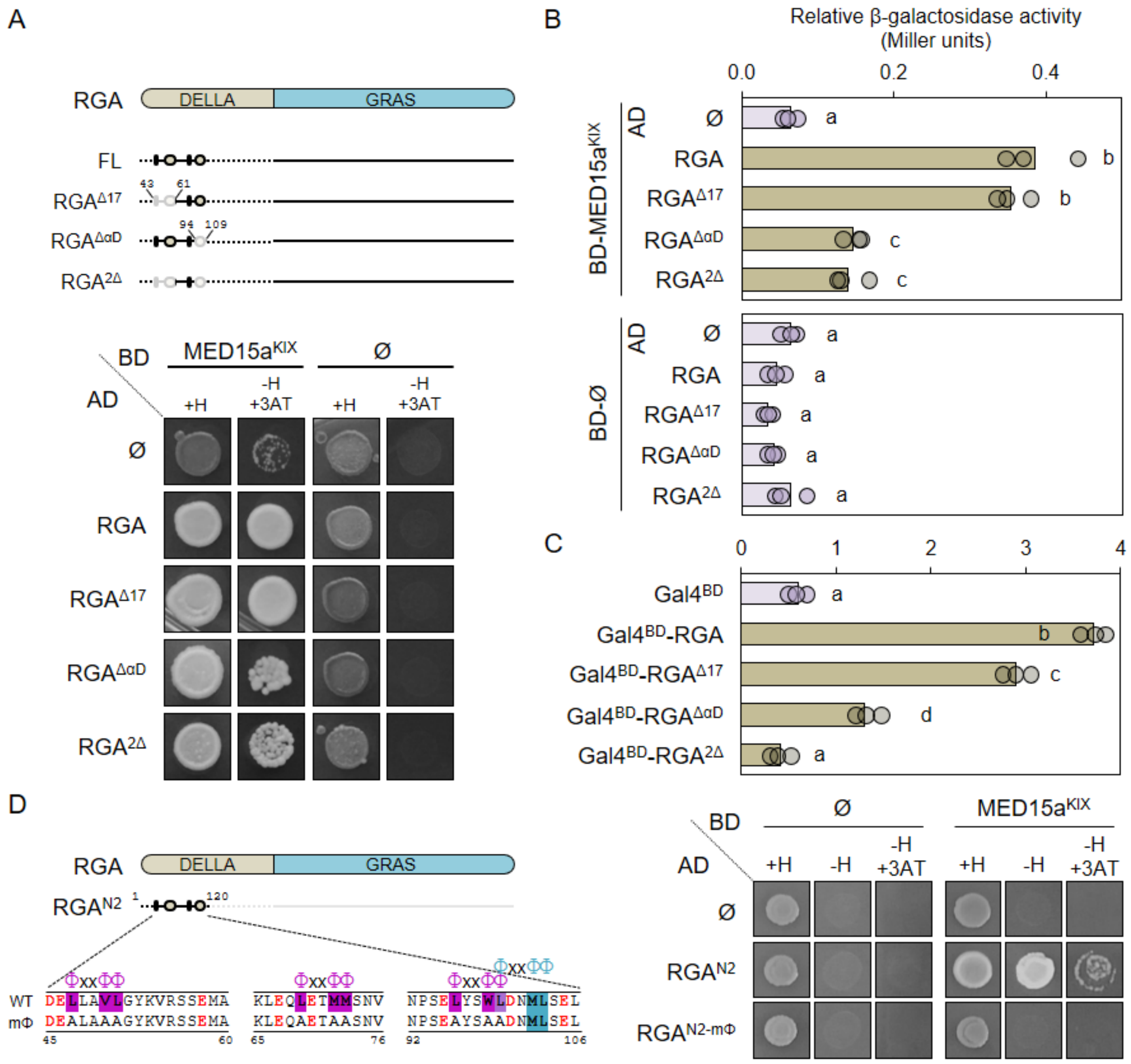


**Fig. S3. p53-like motifs in DELLA domains drive gene activation and MED15 recruitment**. A) Upper panel, illustration of RGA deletions used. MR, Middle Region; MAD, Mediator Activation/Association Domain. $RGA^{2\Delta}$ is a combination of $RGA^{\Delta17}$ and $RGA^{\Delta\alpha D}$ deletions. Ovals represent α helixes based on known structures. Lower panel, yeast two-hybrid drop assay of RGA deletions (fused to the GAL4 activation domain, AD) with the MED15a KIX domain (fused to the GAL4 DNA binding domain, BD). B) Quantification of yeast two-hybrid LacZ reporter activation of the yeast lines in A). C) Yeast one-hybrid of GAL4 DNA binding domain ($Gal4^{BD}$) fused to the RGA deletions depicted in A), shown as the quantification of the LacZ reporter activation. D) Left panel, illustration of the RGA deletion N2 version and its sequence highlighting the p53-like motifs and the mutant version (mΦ). Right panel, yeast two-hybrid drop assay showing the interaction of the N2 region of RGA with MED15a KIX domain. Data in B and C represents the mean activity (bar) of three independent biological replicates (circles).

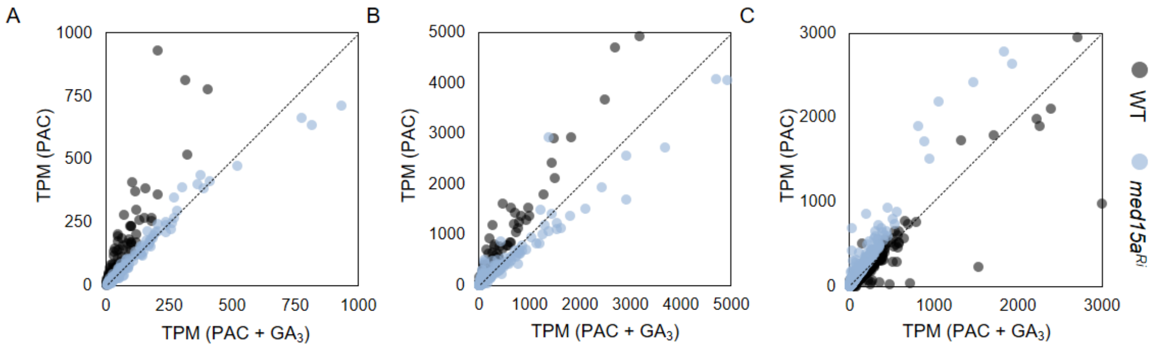


**Fig. S4. DELLA up-regulated genes induction requires MED15**. A) Expression values (as TPMs) of the 184 genes in the intersection between those up-regulated by PAC compared to PAC+$GA_3$ treatment in wild-type (WT, gray) and *med15a$^{Ri}$* line (blue). B, C) Full zoomed-out scatter plot of TPM count represented in Fig. 2B plots.

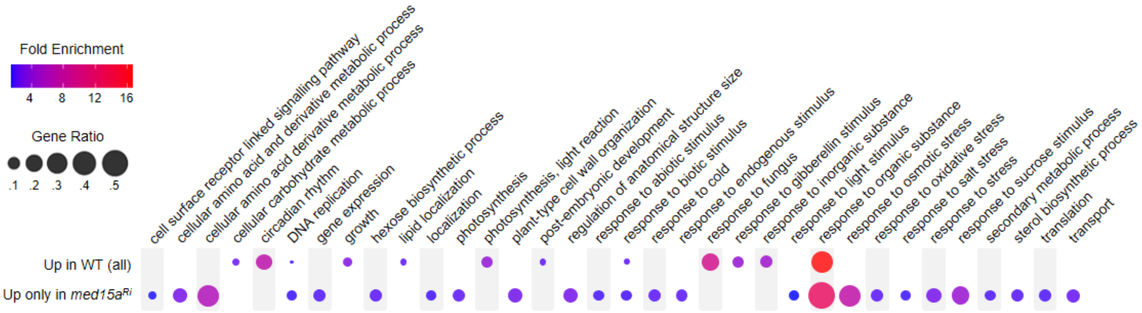


**Fig. S5. Gene ontology analysis of the DELLA up-regulated genes suggest MED15 involvement in gibberellin regulated physiological responses.** GO enrichment only in Biological Processes found for the list of genes up-regulated by PAC in the *MED15a* RNAi line, or all the genes up-regulated by PAC in the wild-type, which is accounted as the list of putative DELLA-MED15 targets. GO enrichment was first calculated based on AgriGO. Code for enrichment plot can be found in the Mendeley Data resource.

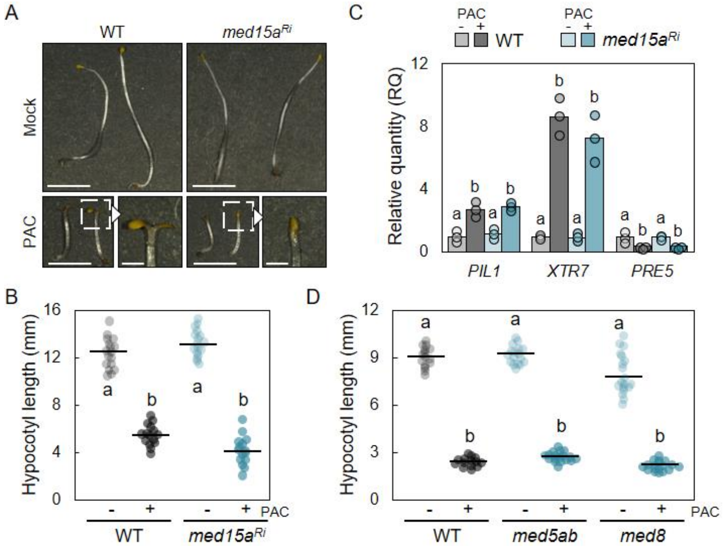


**Fig. S6. Disruption of MED15 function does not affect DELLA inhibition of PIF-mediated responses**. A) 5 days-old wild-type and *med15a$^{Ri}$* seedlings grown in darkness with or without 1 µM PAC. Close-up pictures emphasizing the cotyledon opening phenotype. Scale bars, 5 mm; close-up, 1 mm. B) Quantification of hypocotyl length in A. C) RT-qPCR analysis of GAs and PIFs responsive genes in 5 days-old wild-type and *med15a$^{Ri}$* seedlings grown in darkness with or without 1 µM PAC. Data are medians (bar) of 3 biological replicates, referred against mock and 2 technical repeats (means per replicate shown as filled circles), referred against mock. PDF2.1 was used to normalize data. D) Hypocotyl length of 5 days-old wild-type, *med5ab*, and *med8* seedlings grown in darkness with or without 1 µM PAC. B and D show experimental data of two independent replicates merged with at least 15 plants per genotype and treatment. Dots represent individual plants and horizontal lines total mean values. Statistical groups are determined by Tukey's Post-Hoc test ($p<0.05$) following ANOVA analysis. In C), dots represent biological replicates from three independently performed experiments, and ANOVA analyses performed independently per gene.

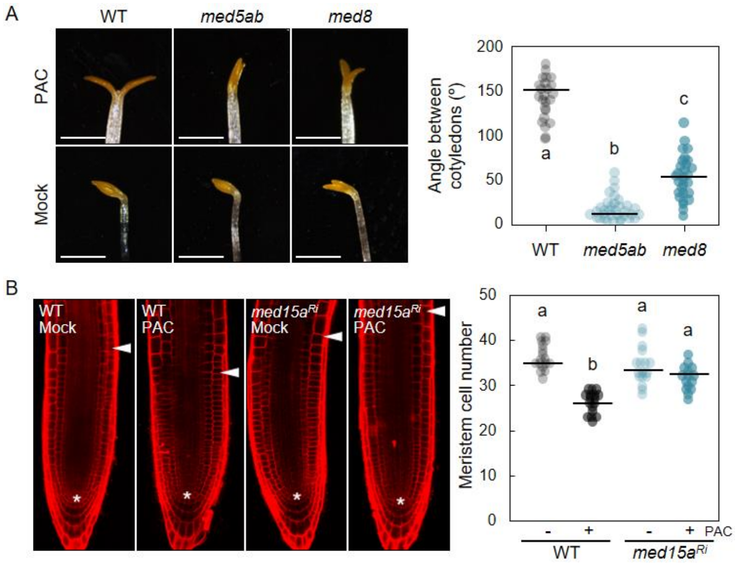


**Fig. S7. Mediator disruption affects DELLA-mediated cotyledon opening and root apical meristem size regulation**. A) Left, 5 days-old WT and *med5ab* and *med8* mutant seedlings grown in darkness with or without 1 µM PAC. Scale bar, 1 mm. Right, dot plot of angle between cotyledons in these lines. Measurements in mock plants not shown (mean = 0; SD = 0). B) Left, confocal microscope images of representative root apical meristems of 5 days-old *med15a*$^{Ri}$ seedlings grown with or without 10 µM PAC for 16 hours. Arrowheads point to the interface between the meristem and the elongation region; asterisks mark the quiescent centers. Right, meristem size measured as cortex cell number. Data represents two merged replicates with at least 10 plants (A) and 7 plants (B) per genotype and treatment. Statistical groups in A and B graphs determined by Tukey's Post-Hoc test ($p<0.05$) following ANOVA analysis.

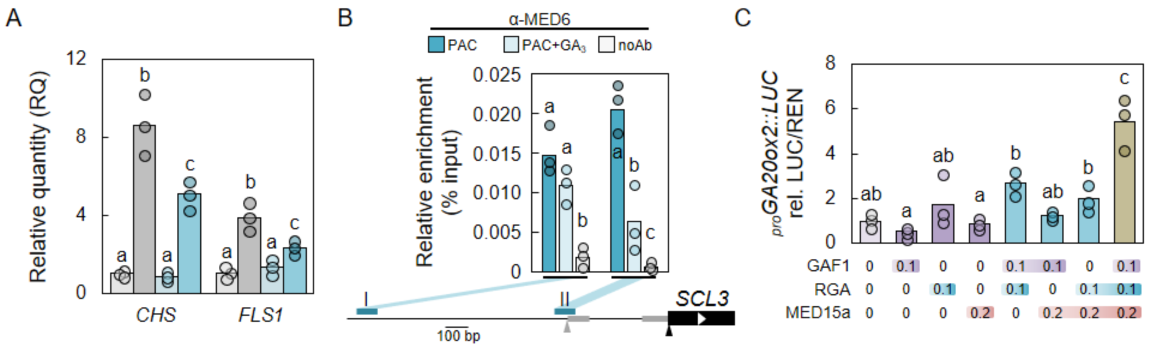


**Fig. S8. Mediator is necessary for DELLA activation of target genes**. A) RT-qPCR analysis of GA and MYB12-responsive genes in 5 days-old WT and *med15a*$^{Ri}$ seedlings grown in darkness with or without 1 µM PAC using *PDF2.1* as reference gene. B) ChIP-qPCR analysis of MED6 occupancy at the indicated locations of *SCL3* promoter in 7 days-old seedlings treated with 1 µM PAC, and with or without 100 µM $GA_3$ for 3 days. Relative enrichment is given as percentage of IP/Input. C) Dual luciferase transactivation assay in *N. benthamiana* leaves using the *LUC* gene under the control of *GA20ox2* promoter as the reporter, and HA-GAF1, YFP-RGA, and cMyc-MED15a co-expressed as effectors. Effector level is indicated below each bar as the agroinfiltrated $OD_{600}$. Bars represent biological replicate medians, and circles represent means of technical triplicates per biological replicate. B) and C) bars are means of 3 biological replicates. Statistical groups determined by Tukey's Post-Hoc test ($p<0.05$) following ANOVA analysis. For qPCR, ANOVA analyses were performed independently per gene/genomic region.

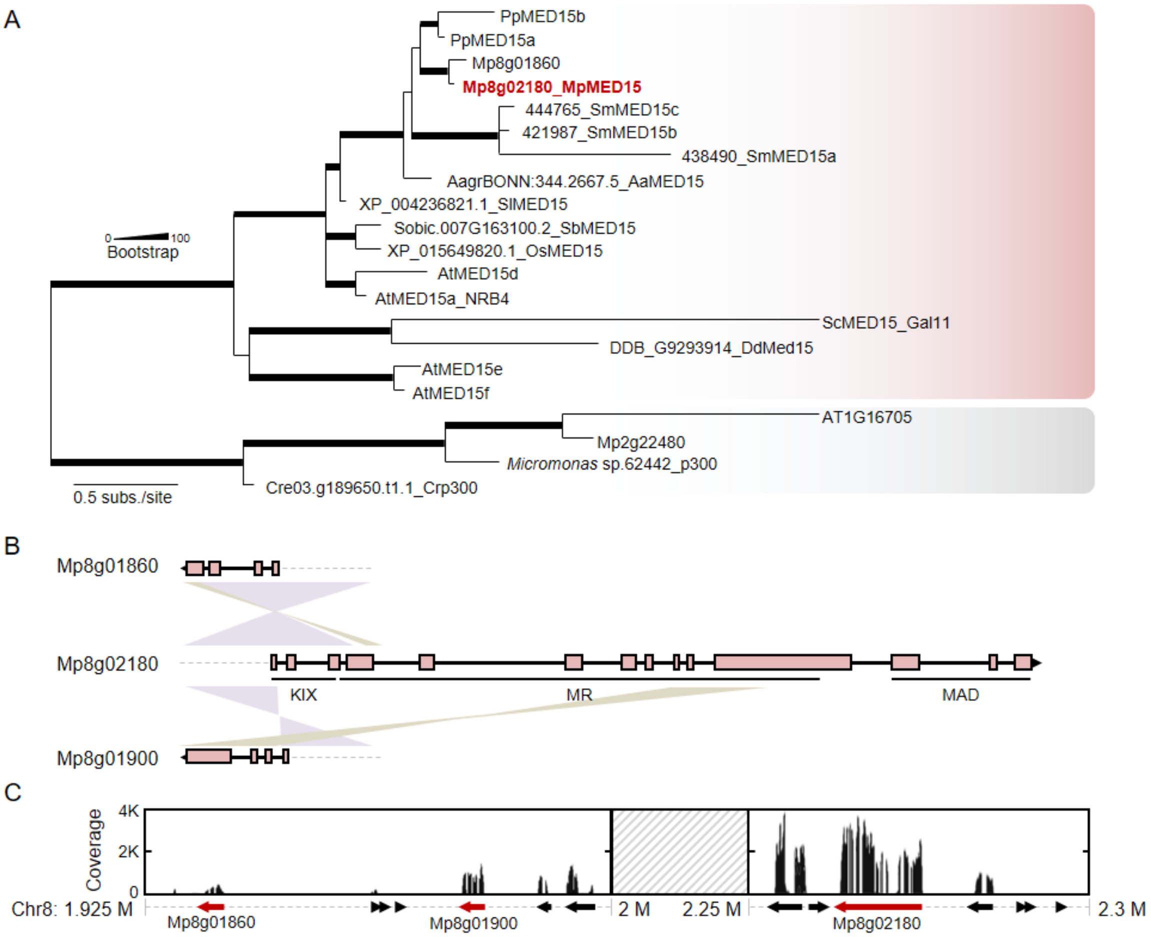


**Fig. S9. Identification of *MED15* genes in *Marchantia polymorpha***. A) Phylogenetic analysis of MED15 proteins using KIX domains. Grey shaded clade is composed of CBP-type KIX domains and represents the outgroup. Support values associated with branches and displayed as bar thickness are maximum likelihood bootstrap values from 1.000 replicates. Branch length represent distance in substitutions per site. B) *MED15* genes *Mp8g01860* and *Mp8g1900* sequence coincidence (>90% identity) with *Mp8g02180* gene. MR, Middle Region; MAD, Mediator Activation/Association Domain. B) RNA-seq coverage of minus (-) strand of *Marchantia polymorpha* Tak-1 wild-type chromosome 8. Red arrows represent *MED15*-related genes, while black arrows are unrelated genes. Data extracted from the *marchantia.info* database.

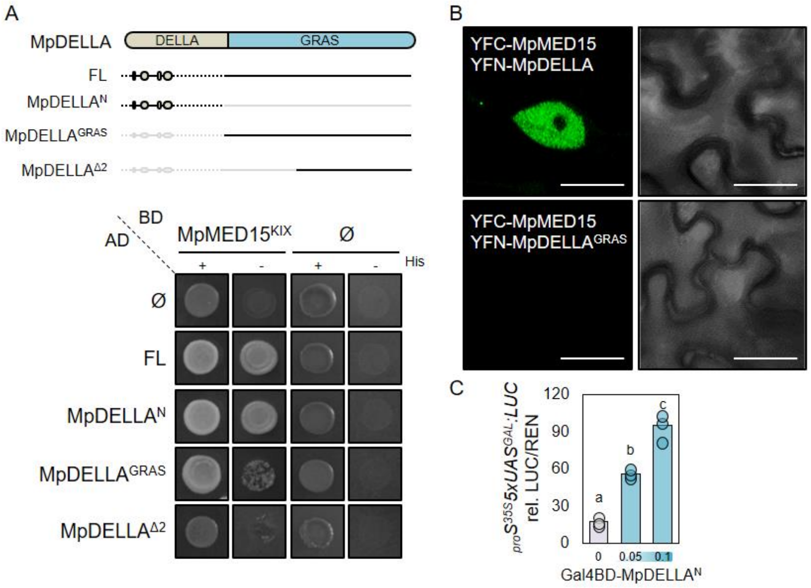


**Fig. S10. MpMED15-MpDELLA interaction analysis.** A) Up, illustration of MpDELLA deletions. Down, yeast two-hybrid assay using MpMED15$^{KIX}$ as bait (Gal4BD), and MpDELLA deletions as prey (Gal4AD). B) Bimolecular fluorescence complementation assay of MpMED15 and MpDELLA, using MpDELLA GRAS domain-only as negative control. C) Dual luciferase using the *LUC* gene under the control of the *Gal* operon UAS promoter as the reporter (5*xUAS$_{Gal}$::LUC*) and the GAL4 DNA binding domain (BD) fused to the MpDELLA DELLA domain (MpDELLA$^{N}$) as effector. Data represent means (bar) of three biological replicates. Circles are mean values of three technical replicates in a biological replicate. Letters indicate statistically different groups calculated by one-way ANOVA with a Tukey HSD post-hoc analysis ($p < 0.01$).

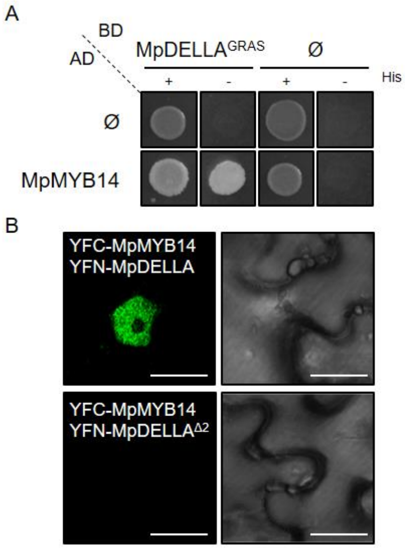


**Fig. S11. MpDELLA-MpMYB14 interaction analysis**. A) Yeast one-hybrid assay using MpDELLA GRAS domain as bait (Gal4BD), and MpMYB14 as prey (Gal4AD). B) Bimolecular fluorescence complementation assay of MpMYB14 and MpDELLA, using a MpDELLA deletion version as negative control. Scale bar, 10 µm.

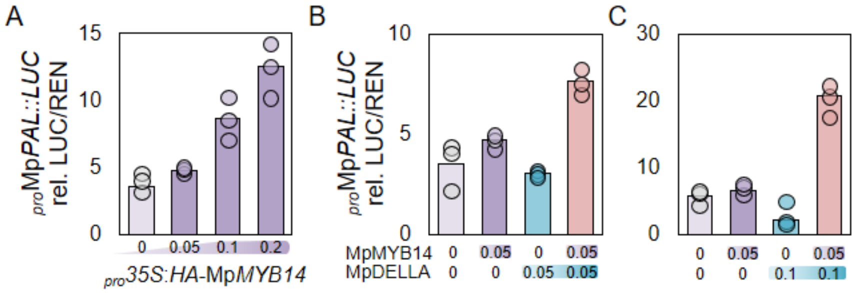


**Fig. S12. MpDELLA-MpMYB14 co-activate a *pro*Mp*PAL*-driven reporter gene.** Dual luciferase using the *LUC* gene under the control of the Mp*PAL* gene promoter, and the HA-fused MpMYB14 as effector alone (A), or together with YFP-fused MpDELLA (B & C). Effectors were infiltrated with a preparation of *A. tumefaciens* with the optical densities (O.D.) indicated below the bar plots. Data represent means (bar) of three biological replicates. Circles are mean values of three technical replicates in a biological replicate. Letters indicate statistically different groups calculated by one-way ANOVA with a Tukey HSD post-hoc analysis ($p < 0.01$).

**Dataset S1 (separate file).** DEG genes in wild-type and med15aRi padj0.01.

**Dataset S2 (separate file).** EAT-UpTF analysis using up-regulated genes in the wild-type subset and in the MED15aRi-only.

**Dataset S3 (separate file).** GO enrichment in DELLA-MED15 common targets and genes up-reg in med15aRi only.

**Dataset S4 (separate file).** Plasmids, primers and protein alignments used in this study.
